# Human-specific fast synaptic kinetics enable rapid detection of predictive features

**DOI:** 10.1101/2025.11.03.686448

**Authors:** Kisho Obi-Nagata, Norimitsu Suzuki, Kenzo Kosugi, Katsuya Ozawa, Hiroki Sasaguri, Kimie Niimi, Kentaro Miyamoto, Masaki Takao, Masahiro Toda, Toshitake Asabuki, Akiko Hayashi-Takagi

## Abstract

Humans can discern the underlying principles in complex environments, but the cellular basis of this ability is unclear. By comparing layer 2/3 cortical pyramidal neurons in five mammals (mouse, rat, marmoset, macaque and human), we found that human neurons exhibited distinctively faster kinetics of excitatory synaptic transmissions. In a computational model, neurons with human-type synapses detected hidden patterns more quickly in noise, and this superiority was only evident in unsupervised learning. Tracking synaptic weights revealed that human synapses were potentiated sharply and persistently by the patterns, while macaque-type synapses were broadly potentiated, which were easily eroded. These data suggest that the human-specific acceleration of synaptic kinetics enables the rapid extraction of latent structures in complex environments, providing a cellular basis for our efficient cognitive abilities.

## Introduction

The human brain is highly adept at swiftly and accurately identifying significant, though often subtle, features in noisy environments. This ability is essential for making quick and appropriate decisions. Substantial efforts have been made to identify the biological mechanisms that account for the superiority of human brain function. Structurally, the human neocortex is larger, contains more neurons, and exhibits more elaborate neurites with more abundant synapses compared to other mammals (*1-3*). Functionally, electrophysiological studies using human acute cortical slices resected from patients undergoing surgery have demonstrated that human neurons are not merely scaled-up versions of neurons found in other species. Instead, there are properties specific to humans that include distinct passive and active membrane characteristics, as well as action potential dynamics (*2, 4-12*). Despite these advances, it remains unclear which neuronal features underlie the mechanisms enabling higher-order cognitive functions. Two limitations that have impeded progress. First, despite the pivotal role of the neocortex in human cognition, comparative work in the neocortex has centered on humans and mice, leaving it uncertain whether prior findings are uniquely human or broadly primate. Second, there has been a paucity of functional and quantitative analyses of human synapses in the neocortex. Although recent progress in structural connectomics using electron microscopy (*3*) and functional connectomics by multicellular patch-clamp recordings (*13-16*) has provided fine wiring maps, these approaches lack information on functional contributions of individual synapses on neural computation. Theoretical work suggests that increasing the number of neurons or synapses improves performance only up to a point. However, beyond this threshold, which is far less than that of the human brain, the introduction of additional connections amplifies noise and degrades performance (*17*). In addition, given the highly non-linear nature of synaptic integration (*18-21*), it is imperative to undertake not only the enumeration but also the quantitative assessment of individual synaptic properties and subsequent firing in order to comprehend human cortical computation. Here, we scrutinized excitatory postsynaptic currents and potentials (EPSCs/EPSPs) in layer 2/3 pyramidal neurons from acute slices of dorsolateral prefrontal and temporal cortex obtained from human patients. We compared them to those from mouse, rat, marmoset, and macaque under identical experimental conditions. The particular focus is on scrutinizing two non-human primates, such as marmosets and macaque monkeys, and comparing them to humans. We focused on layer 2/3, because this layer expanded disproportionately in humans (*22*), forming the cortico-cortical recurrent circuits for the superiority of human flexible cognition. Furthermore, to examine their functional consequences, we used a computational model endowed with either human or macaque synapses to reveal the computational advantage of fast kinetics in pattern detection.

## Results

### Passive membrane properties of neurons from five mammalian species

To identify the human-specific physiological characteristics of cortical pyramidal neurons, we prepared acute brain slices from the prefrontal or temporal cortex of five mammalian species (mouse, rat, marmoset, macaque, and human) under practically identical experimental conditions. Human cerebral cortex samples were obtained from the brains of patients undergoing surgery for therapeutic purposes for glioma, metastatic tumor, or epilepsy (Table 1). The preoperative MRI scan and pathological examination showed that the resected areas were not pathological, with no signs of cancer cell infiltration or other neurological abnormalities (fig. S1). We focused on layer 2/3 because the human neocortex is heterogeneous across layers (*15, 22*), and superficial layer 2/3 pyramidal neurons are a relatively homologous cell type with similar morphological features – dendritic length, somatic size, and cortical depth – across mammalian species (Fig. 1A, fig. S2). Neurons were classified as excitatory based on their morphology and intrinsic electrophysiological properties (Fig. 1A, B). We first quantify somatic passive membrane parameters and firing characteristics, including resting membrane potential, input resistance, membrane time constant, action potential threshold, and rheobase. These properties were relatively conserved across species, with some significant differences observed: human neurons had a shorter membrane time constant than rodent and marmoset neurons, and an enhanced sag ratio compared to their rodent counterparts (Fig. 1B–M). Overall, these somatic membrane properties are consistent with previous findings (*6*).

**Fig. 1.**
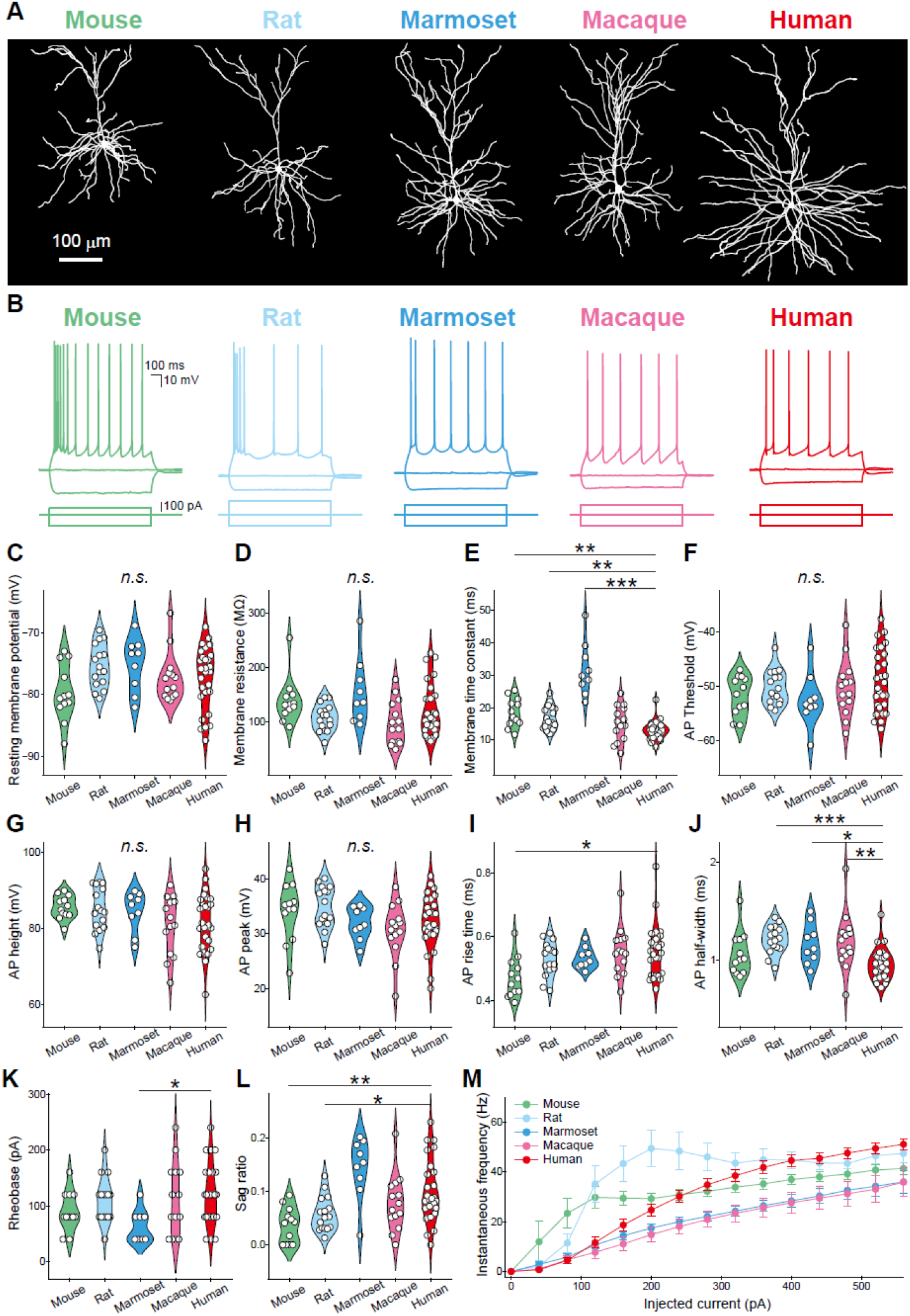
Passive membrane properties of neurons from five mammalian species. (**A, B**) Representative images of the layer 2/3 cortical neurons analyzed using whole-cell patch-clamp (**A**) and somatic voltage responses to step current injections (**B**). (**C-M**) Somatic membrane properties. Resting membrane potential (**C**). Membrane resistance (**D**). Membrane time constant (**E**). Action potential threshold (**F**). Action potential height (**G**). Action potential peak (**H**). Action potential rise time (**I**). Action half-width (**J**). Rheobase (**K**). Sag ratio (**L**) The firing rates are depicted as a function of the injected current (**M**). Error bars based on the standard error of the mean (s.e.m.). Mouse, *n* = 12 cells; Rat, *n* = 16 cells; Marmoset, *n* = 9 cells; Macaque, *n* = 14 cells; Human, *n* = 28 cells. \**P* < 0.05, \*\**P* < 0.01, \*\*\**P* < 0.001. See also **Table S2** for measurements and statics.

### Human-specific fast kinetics of excitatory postsynaptic events

We next compared spontaneous EPSCs (sEPSCs) across species, analyzing neurons matched for soma size and cortical depth (Fig. 1A, fig. S2). The most striking characteristic of human sEPSCs was the sharp kinetics, manifested as both faster rise and decay times (Fig. 2B, C, E, F). These steeper kinetics were specific to humans, whereas those observed in non-human primates, such as macaques and marmosets, were slower and comparable to those observed in rodents (Fig. 2H, I). As previously reported (*12*), humans have a notably larger amplitude of sEPSCs than rodents, and the amplitude of human sEPSCs was identical to that observed in marmosets and macaques (Fig. 2D, G, J). We then investigated whether human EPSPs are faster than those of other species. This is an important consideration because EPSPs directly contribute to action potential generation and their values are largely influenced by geometric factors, such as synapse-to-soma distance and dendrite thickness (*2*). Measuring sEPSP kinetics revealed that the rise and decay times of humans were significantly shorter than those of other animals (Fig. 3). Furthermore, to directly assess the relationship between EPSPs and these geometric factors, we employed two-photon glutamate uncaging of 4-methoxy-7-nitroindolinyl glutamate (MNI-Glu) at single spines to precisely stimulate the dendritic spines under patch-clamp conditions and recorded the uncaging-evoked excitatory postsynaptic potentials (uEPSPs) (Fig. 4). The rise time of uEPSPs in humans was significantly shorter than in the other four species (Fig. 4C). In terms of decay time, no significant difference was noted between humans and macaques; however, the decay time of human uEPSPs was markedly shorter than that of mice, rats, and marmosets (Fig. 4D). Lastly, we asked whether the observed phenotypes in human neurons were specific to humans or could be explained by confounding factors. To investigate this, we examined the effects of transcardiac perfusion with a PBS solution prior to resection, synapse-to-soma distance, and donor age (fig. S2–5). The presence or absence of transcardiac perfusion had no effect on EPSC kinetics (fig. S3). There was no correlation between the location of the recorded cells in the cortex and their relationship to EPSC kinetics (fig. 2). Furthermore, precise identification of the synaptic location via single-spine stimulation using two-photon glutamate uncaging showed no consistent correlation between synapse-to-soma distance and uEPSC kinetics (fig. S4). Interestingly, we observed a significant negative correlation between the donor age and EPSC time course in marmosets and humans (fig. S5). Taken together, the faster kinetics in humans synaptic transmission reflect intrinsic synaptic properties rather than other confounding factors.

**Fig. 2.**
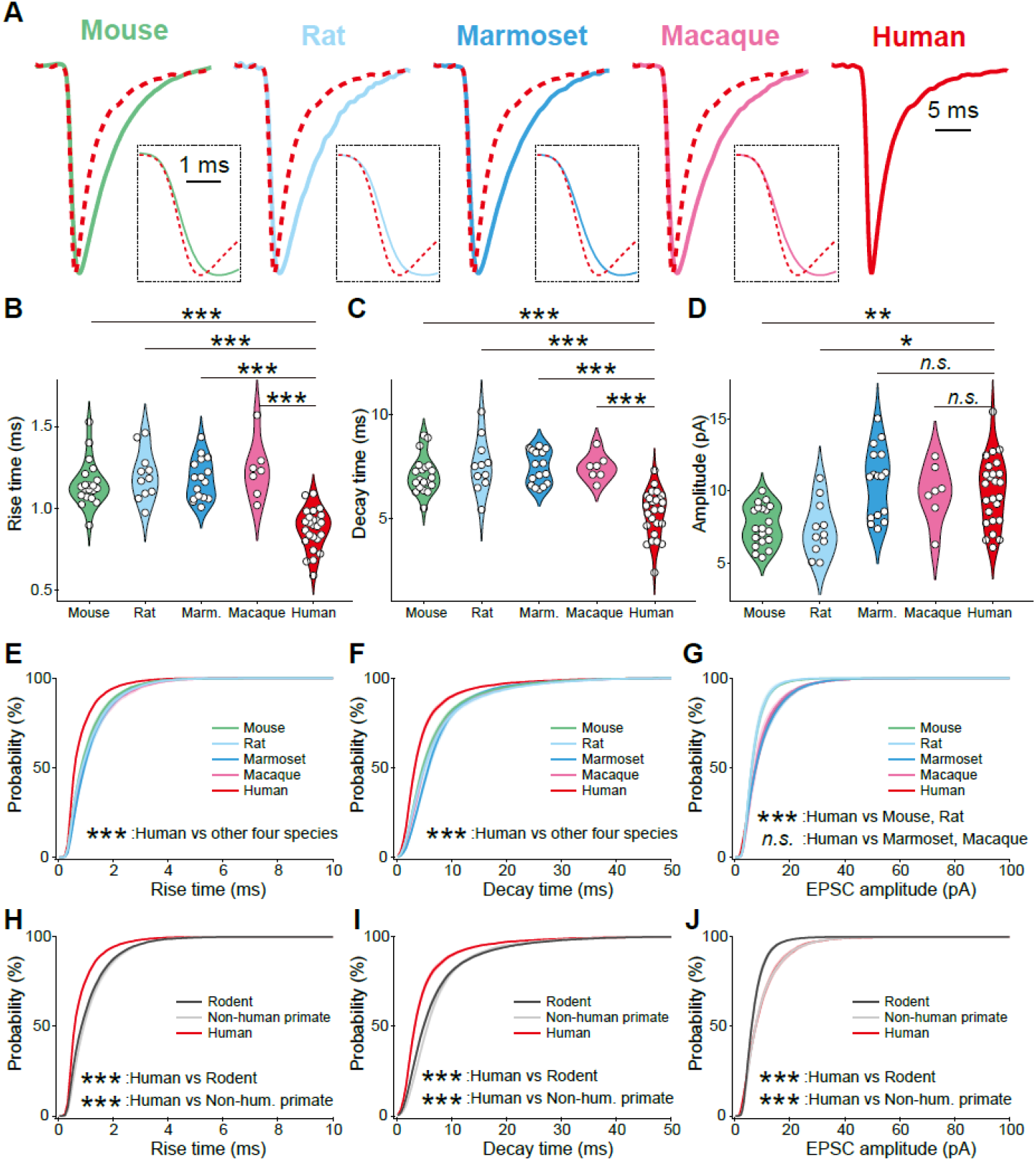
Human-specific features of excitatory postsynaptic events. (**A**) Representative EPSCs from five species. The human EPSC trace is superimposed as a red dotted line for comparison. Inset: higher magnification of the rising phase of the EPSCs. (**B**) Violin plots showing rise time, as measured between 20% and 80% of the EPSC amplitude. (**C**) Violin plots showing the decay time of EPSCs, extracted by a single exponential decay fitting. (**D**) Violin plots showing the amplitude of EPSCs. (**E–J**) Cumulative density plots showing the rise time, decay time, and amplitude of EPSCs for the comparison of the five species (**E–G**) and that of rodent, non-human primate (marmoset and macaque), and human (**H–J**). Mouse, *n* = 20 cells; Rat, *n* = 11 cells; Marmoset, *n* = 20 cells; Macaque, *n* = 7 cells; Human, *n* = 25 cells. \**P* < 0.05, \*\**P* < 0.01, \*\*\**P* < 0.001. See also **Table S2** for measurements and statics.

**Fig. 3.**
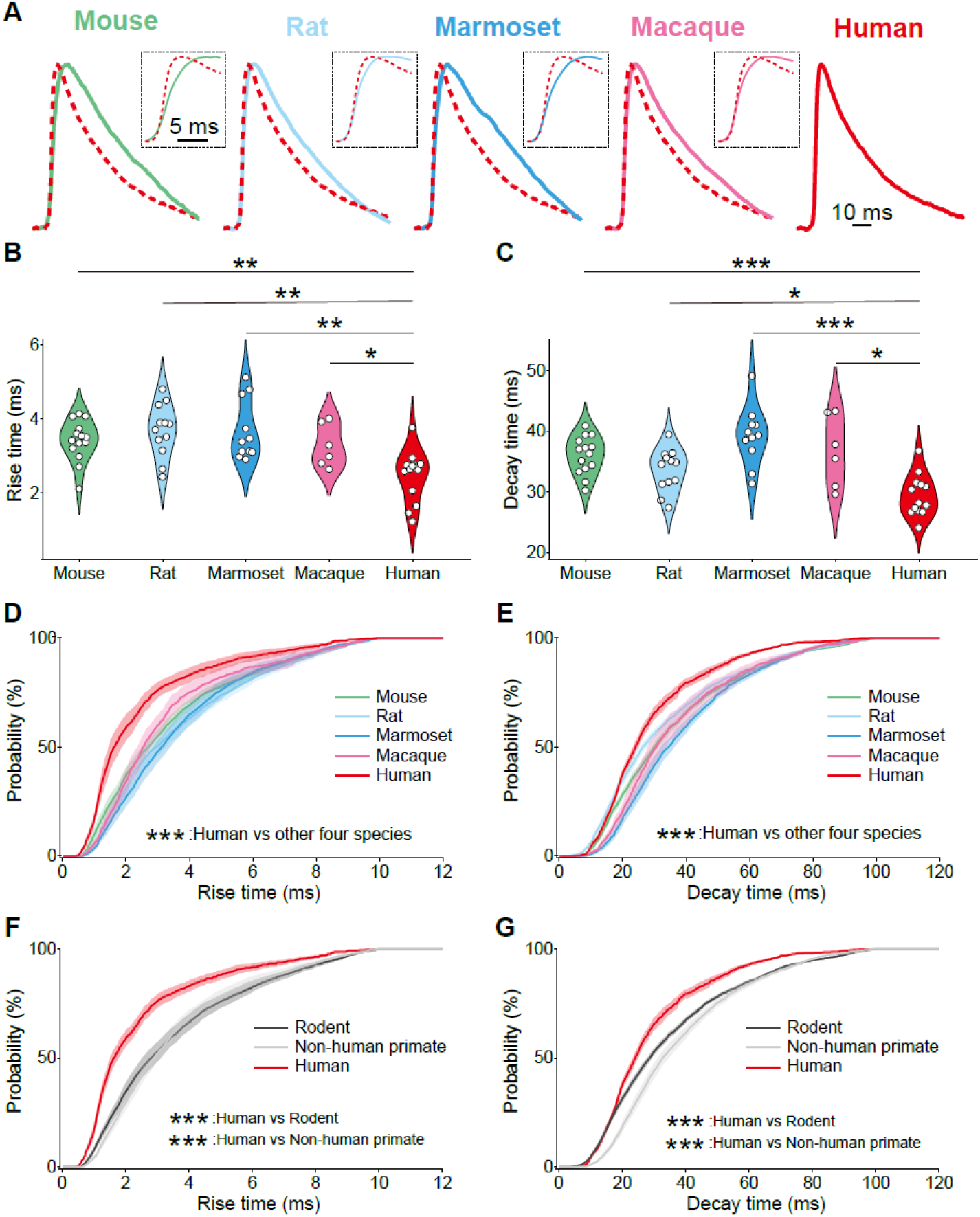
Human-specific fast kinetics of EPSPs. (**A**) Representative trace of the EPSPs. The human EPSP trace is superimposed as a red dotted line for comparison. Inset: higher magnification of the rising phase of the EPSPs. (**B, C**) Violin plots showing rise time (**B**) and the decay time (**C**) of EPSPs. (**D, E**) Cumulative density plots showing the rise time, decay time, and amplitude of EPSPs for the comparison of the five species (**F, G**) and that of rodent, non-human primate (marmoset and macaque), and human (**H–J**). Mouse, *n* = 14 cells; Rat, *n* = 12 cells; Marmoset, *n* = 11 cells; Macaque, *n* = 6 cells; Human, *n* = 13 cells. \**P* < 0.05, \*\**P* < 0.01, \*\*\**P* < 0.001. See also **Table S2** for measurements and statics.

**Fig. 4.**
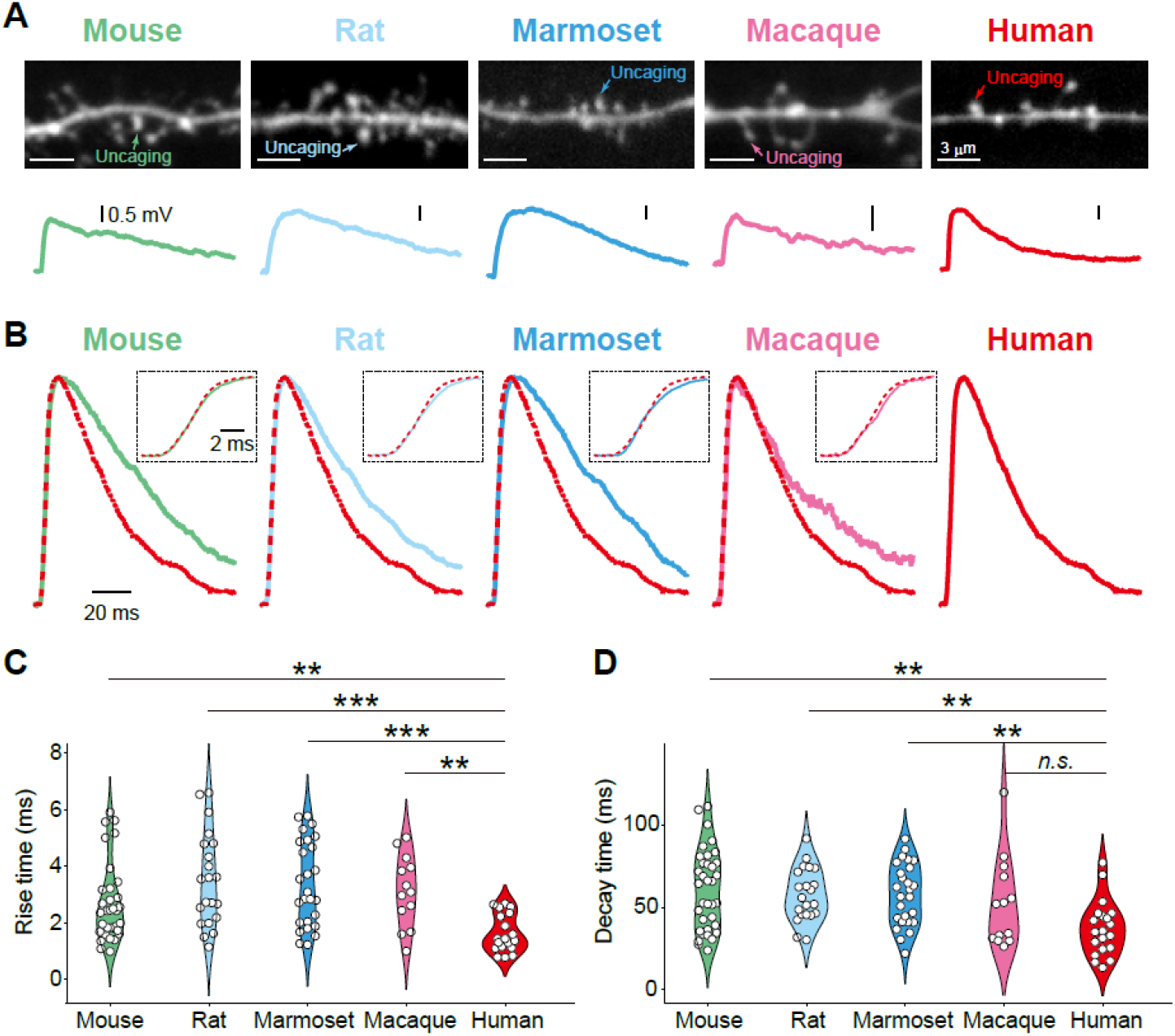
The sharp kinetics of human EPSPs evoked by single-spine stimulation. (**A**) A dendritic spine subjected to single-spine stimulation by glutamate uncaging and the corresponding EPSP upon uncaging (uEPSP). (**B**) uEPSPs trances from five species. The human uEPSP trace is superimposed as a red dotted line for comparison. Inset: higher magnification of the rising phase of the uEPSPs. (**C, D**) The rise time (**C**) and decay time (**D**) of uEPSPs. Mouse, *n* = 35 spines; Rat, *n* = 21 spines; Marmoset, *n* = 26 spines; Macaque, *n* = 13 spines; Human, *n* = 20 spines. \*\**P* < 0.01, \*\*\**P* < 0.001. See also **Table S2** for measurements and statics.

### Fast synaptic kinetics enhance unsupervised learning of predictive features

We next tested whether and how human-specific fast synaptic kinetics affect sequence learning in a two-compartment neuron model (Fig. 5A). In this model (*23*), dendrites receive input and generate local responses, while the soma produces outputs. Synaptic weights are updated to match dendritic and somatic responses over short histories, with adaptive modulation of gain and threshold preventing trivial solutions. We compared neuron models with distinct EPSP kinetics: fast-rise/fast-decay (human/human type, hereafter “human-type”), slow-rise/slow-decay (macaque/macaque type, hereafter “macaque-type”), and two artificial chimeric forms (slow-rise/fast-decay and fast-rise/slow-decay, macaque/human type and human macaque type, respectively) constrained by experimental measurements. In addition, we included a mouse-type neuron defined by experimentally measured mouse EPSP kinetics (slow-rise/slow-decay, distinct from macaque values). The model received Poisson-like input from a large presynaptic population, with 10-ms spatiotemporal patterns intermittently embedded in irregular spike trains (Fig. 5B, top), similar to previous modeling studies (*24, 25*). These patterns could not be inferred from mean firing rate alone. When trained on the input stream, the human-type neuron developed selective responses to the recurring spike pattern, indicating that the model successfully learned the embedded temporal structure without external supervision (Fig. 5B, bottom). We next examined how the learned behavior differed between the models corresponding to two species (*i*.*e*., human- and macaque-type). To this end, we first quantified the synaptic weight changes associated with neurons participating in the embedded pattern. In both human- and macaque-type models, synaptic connections from the pattern-participating presynaptic neurons were selectively strengthened during learning, whereas non-participating inputs showed little change (Fig. 5C). Notably, the magnitude of potentiation was significantly greater in the human-type synapses than in the macaque-type synapses, indicating enhanced sensitivity to temporally structured inputs.

**Fig. 5.**
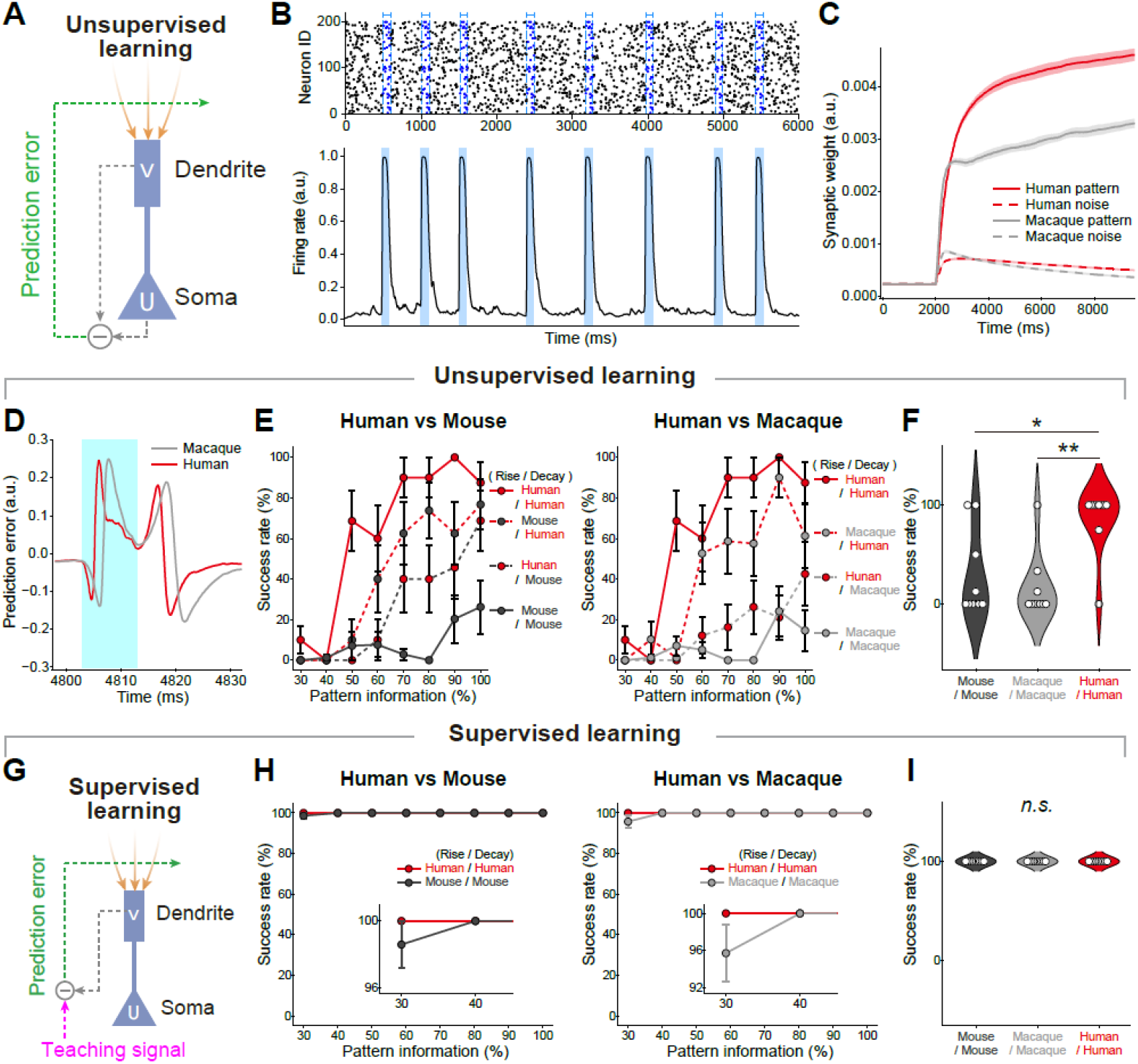
Unsupervised or supervised learning in two-compartment neuron model. (**A**) The model neuron that undergoes unsupervised learning. The dendritic compartment receives input spike trains, and the somatic membrane potential is computed. The somatic output backpropagates to the dendritic synapses as a self-teaching signal. Learning proceeds until the dendrite minimizes the error between its prediction and the actual somatic firing rate. (**B**) Fixed spatiotemporal pattern was embedded repeatedly within irregular spike trains from 200 input neurons. All neurons fire with the firing rate of 5 Hz (Top). The two-compartment neuron learned the recurring patterns. The somatic firing rate before and after learning are shown (Bottom). (**C**) Synaptic weights from pattern-participating inputs increased during learning in both human- and macaque-type models, but the potentiation was greater in the human-type synapse. (**D**) Somato-dendritic prediction error signals were phase-locked to pattern presentations (the period displayed in light blue) in both models, but the responses were sharper and more temporally aligned in the neuron with human-type synapses. (**E**) Degrading the input patterns reduced success rates in all models, yet the human-type neuron remained more robust, showing a gradual performance decline compared with the macaque- and mouse-type neurons. (**F**) Average success rates across models. The human-type neuron consistently outperformed macaque- and mouse-type models. (**G**) The model neuron undergoes supervised learning. The dendritic compartment receives input spike trains, and synaptic updates are driven by the difference between dendritic predictions and external teaching signals. (**H**) Same as (**E**), but for the supervised learning model. (**I**) Same as (**F**), but for the supervised learning model. \**P* < 0.05, \*\**P* < 0.01.

The enhanced potentiation in the human-type model suggests that its distinct synaptic kinetics may facilitate more precise credit assignment during learning. To understand the underlying mechanism, we compared the somato-dendritic prediction error over these models (*26*). We found that although both human and macaque models exhibited prediction error signals phase-locked to the pattern presentations, the human-type neuron showed much sharper responses tightly aligned with the pattern boundaries (Fig. 5D). Specifically, human-type kinetics generated narrow negative-to-positive peaks at pattern onset and positive-to-negative peaks at offset, precisely marking the boundaries. In contrast, macaque-type and mouse-type kinetics produced broader, delayed peaks, extending the effective window of synaptic plasticity and reducing the temporal precision of boundary encoding. This sharpening arose because fast EPSP kinetics restricted the dendrite’s integration window, enabling precise spatiotemporal detection of boundaries between the pattern and noise.

We next tested robustness by degrading patterns through random spike omissions (30–100% completeness) to mimic real-world environments in which animals must learn amidst insufficient information. We found that performance of all models declined as patterns were degraded, with outcomes strongly dependent on kinetics (Fig. 5E, F). Human-type neurons consistently outperformed all others across completeness levels and showed the slowest performance decline. Mouse-type and macaque-type neurons performed worse, with low accuracy even for intact patterns and a steep deterioration under degradation. The performance of the chimeric synapses was intermediate, with decay speed more influential than rise (Fig. 5E, Cohen’s d = 0.45 for rise vs. 0.79 for decay). Interestingly, in supervised learning model, all synapses achieved comparable performance (Fig.5G–I), suggesting that kinetic differences become critical only when learning relies on internally generated teaching signals. Altogether, these results suggest that fast human-type kinetics may support more efficient internal credit assignment during unsupervised learning.

## Discussion

Our understanding of human synapses, especially compared to non-human primates, is limited. This is problematic because studies in rodents have shown that circuit algorithms reside in synapses, and that synaptic weights and kinetics markedly affect neural computation — a finding that should also apply to the human brain. For example, spike-timing-dependent plasticity (*27*), behavioral time scale synaptic plasticity (*28*), and or gating and integration of multiple sources (*29*) are largely dependent on synaptic and resultant dendritic/somatic events. We here scrutinized sEPSC, sEPSP, and uEPSP of pyramidal neurons in the superficial layers 2/3 neocortex. An advantage of focusing on layers 2/3 is that the length and thickness of dendrites, as well as somatic size, vary less between species than in deeper layers. Since these neuronal morphologies influence function, standardizing conditions other than synapses allows for a more rigorous comparison of synaptic function across species. This is not only the first in-depth functional analysis of human synapses in the neocortex, but also provides a detailed comparison between humans and two non-human primates — macaques and marmosets — to identify characteristics unique to humans. We concluded that fast EPSC/EPSP kinetics were indeed unique to humans, even when various confounding factors were taken into account.

The computational model suggested that neurons with human-type synapses rapidly detect hidden patterns, particularly when information was incomplete or noisy. Chimeric synapses, where the rise and decay times were a combination of macaque and/or human, showed that contributions from both parameters were critical, with decay slightly more influential. At first, faster synaptic kinetics seemed disadvantageous for synaptic plasticity, because temporal summation of faster synaptic events is more difficult, which makes the synaptic plasticity less likely. However, monitoring changes in synaptic weight during learning revealed that potentiation in human-type synapses was finely tuned to specific patterns and was more resilient to depotentiation. These findings suggest that sharp EPSPs may enable rapid and localized credit assignment, allowing dendrites to match top-down predictions with bottom-up inputs on a fine temporal scale. In this view, human-type synapses support not merely Hebbian coincidence detection, but predictive learning based on temporally precise error signals within local dendritic compartments.

In contrast, the synaptic kinetics in the human hippocampal CA3 region are the opposite, with a longer integration time that greatly enhances memory storage capacity (*30*). The difference between these two brain regions highlights the distinct functions of the hippocampus—storing memories (*31*)—and the neocortex—filtering and integrating external stimuli with internal models to form the decision of animals. Interestingly, human synapses in the neocortex only demonstrated superiority under unsupervised conditions, suggesting a link to functions such as intuition and epiphany. Herbert A. Simon said that ‘intuition is nothing more and nothing less than recognition’, describing that intuition isn’t coincidence, but rather an unconscious ability to recognize patterns, providing quick and accurate answers without deliberate thought (*32*). Interestingly, our data showed that steeper EPSCs were more prevalent in older individuals. While many fluid abilities decline with age, some research showed that crystallized/semantic knowledge, elements of wise social reasoning often improve or are preserved (*33-35*). Thus, synaptic features associated with faster integration could still contribute to intuitive, knowledge-driven inferences in later life.

An open question is the molecular mechanism underlying the fast kinetics of human synapses. The existence of genetic architecture that is unique to humans has also recently attracted attention (*36-42*). AMPAR subtype differences — including subunit composition, splice‐isoforms, and auxiliary proteins — are a highly plausible contributors, but presynaptic and glial components such as glutamate uptake dynamics and multivesicular release synchrony may also play roles (*43*). Further research using super-resolution imaging and electron microscopy analysis, as well as patch-seq analysis, is needed to establish a link between the nanostructure of synapses and their function. Another open question concerns consequences of faster synaptic kinetics at higher scale, such as behavior. Although we provided an in-depth characterization of human synaptic kinetics and the resulting cellular response, the impact of these on circuits and behavior, remains unclear. A previous study of human cortical slices reported that human neocortical circuits demonstrate unique network activity (*12, 13*), suggesting that incorporating human EPSPs into recurrent network models will be an important next step. Such models could test whether fast synaptic kinetics indeed facilitate dendritic credit assignment for predictive learning, thereby bridging molecular and algorithmic levels of neural computation.

Together, our results suggest a human-specific synaptic mechanism that accelerates cortical computation and supports unsupervised learning, providing a mechanistic link between neuronal dynamics and cognitive flexibility. This understanding will inspire the next generation of artificial intelligence, including energy-efficient neuromorphic systems that offer a new conceptual and practical link between biological intuition and computational efficiency.

