## Supplementary material for "Human-specific fast synaptic kinetics enable rapid detection of predictive features": Supple figure and table

Science
AAAS

### Supplementary Materials for

#### Human-specific fast synaptic kinetics enable rapid detection of predictive features

**Authors:** Kisho Obi-Nagata[1,2]†, Norimitsu Suzuki[1]†, Kenzo Kosugi[3]†, Katsuya Ozawa[1], Hiroki Sasaguri[4], Kimie Niimi[5], Kentaro Miyamoto[6], Masaki Takao[7], Masahiro Toda[3], Toshitake Asabuki[8]*, and Akiko Hayashi-Takagi[1]*

**The PDF file includes:**

Materials and Methods
Tables S1, 2
Figs. S1 to S5

#### Materials and Methods
##### Ethical considerations
All animal experiments were approved by the Animal Care and Use Committee of RIKEN. All human experiments were approved by the Wako Human Research Ethics Committee of RIKEN and the Keio University School of Medicine Ethics Committee (approval no. 20221203). Patients consented to the surgical procedures, and a portion of the resected tissue was designated as discarded tissue. Under our approved protocol, such discarded tissue was available for this specific research project without necessitating explicit patient consent. Non-essential samples were extracted by the supervising neuropathologist in accordance with the protocol.

##### Human
Human brain specimens were procured through a collaborative effort with the Department of Neurosurgery at Keio University Hospital. Samples were collected from patients undergoing craniotomy for brain tumor resection or drug-resistant epilepsy. The patient cohort comprised individuals of both sexes, aged between 32 and 72 years (see Table 1 for details). The tumor types included glioma and metastatic brain tumors. Samples were obtained from the frontal, parietal and temporal lobes of the brain. From a neurosurgical perspective, all cortical specimens were collected during clinically indicated resections. Non-pathological regions were identified using intraoperative neuronavigation and visual inspection in order to avoid areas with apparent structural abnormalities. For tumor cases, samples were taken from the cortex adjacent to the lesion and confirmed as structurally and histologically normal through both intraoperative assessment and subsequent pathological examination. In the case of epilepsy, tissue was obtained from the lateral temporal cortex during a standard anterior temporal lobectomy for medically intractable mesial temporal lobe epilepsy. The sampled area showed no epileptiform discharges during preoperative or intraoperative electrocorticography. The resected specimens were promptly placed in an ice-cold, oxygenated *N*-methyl-D-glucamine (NMDG) solution (92 mM NMDG, 2.5 mM KCl, 1.25 mM $NaH_2PO_4$, 30 mM $NaHCO_3$, 20 mM HEPES, 25 mM glucose, 2 mM thiourea, 5 mM ascorbic acid, 3 mM Na-pyruvate, 0.5 mM $CaCl_2$, and 10 mM $MgCl_2$). They were then bubbled with a gas mixture of 95% $O_2$ and 5% $CO_2$, kept on ice and transported by car to RIKEN. This journey took approximately 50 minutes.

##### Animals
C57BL/6J mice and Sprague-Dawley rats were purchased from Japan SLC, Inc. (Japan). The male mice were housed in groups of up to five or individually, and were used for experiments at ages ranging from nine to 40 weeks. Marmosets (*Callithrix jacchus*) were obtained from a breeding colony at the RIKEN Center for Brain Science. They were housed and used when they were between 4–10 years old. The brains of Japanese macaques (*Macaca fuscata*) and rhesus macaques (*Macaca mulatta*) were obtained from animals aged 8–10 years that were euthanised in accordance with the approved schedule once the experiments had been completed. The animals were maintained on a 12-hour dark–light cycle in a controlled environment with a temperature of 23–25 °C for rodents and 27–29 °C for marmosets.

##### Acute slice preparation
***Rodent***: Coronal slices (300 μm thick) were prepared from prefrontal cortex and parietal cortex (somatosensory cortex) as described previously (*44*). In brief, animals were deeply anesthetized and transcardially perfused with ice-cold NMDG solution (in mM) (92 NMDG, 2.5 KCl, 1.25 $NaH_2PO_4$, 30 $NaHCO_3$, 20 HEPES, 25 glucose, 2 thiourea, 5 ascorbic acid, 3 Na-pyruvate, 0.5 $CaCl_2$, and 10 $MgCl_2$. Slices were cut in ice-cold NMDG aCSF with a Leica VT1200S vibratome (Wezlar, Germany), and then transferred to a holding chamber containing NMDG aCSF at 35°C. After 10 min of incubation, the slices were then transferred into a recovery chamber containing HEPES with aCSF (in mM) (92 NaCl, 2.5 KCl, 1.25 $NaH_2PO_4$, 30 $NaHCO_3$, 20 HEPES, 25 glucose, 2 thiourea, 5 ascorbic acid, 3 Na-pyruvate, 2 $CaCl_2$, and 2 $MgCl_2$). All solutions were continuously bubbled with 95% $O_2$ and 5% $CO_2$.

***Non-human primate:*** F Following confirmation of cardiac arrest in both marmosets and macaques, a surgical incision was made in the thoracic cavity and an ice-cold, oxygenated NMDG solution was perfused into the left ventricle while blood was drained from the right ventricle. The skull was then secured in a stereotaxic frame and the cranial bone was removed with a circular diamond cutter to allow easier extraction of the brain. For some marmosets and macaques, the procedure was modified to omit blood drainage. In these cases, the cranial bone was removed directly after cardiac arrest and the brain extracted (non-perfusion condition in fig. S3). The extracted brain was then cut into small pieces in an ice-cold NMDG solution. Coronal slices (300 µm thick) were prepared from the prefrontal and parietal (somatosensory) cortices, in the same way as for the rodent preparation.

***Human:*** Upon arrival at RIKEN, the human specimens were swiftly transferred to an ice-cold NMDG solution. The pia mater and large blood vessels on the surface of the brain were carefully removed. The specimens were then cut into small pieces and sliced to a thickness of 350 µm, in the same way as for rodent preparation.

**Whole-cell patch-clamp recording**

Whole cell Patch-clamp recording was performed from the layer 2/3 pyramidal cell using a MultiClamp 700B amplifier (Molecular Devices, San Jose, CA). Slices were continuously superfused at a rate of 2–3 ml/min with aCSF composed of the following concentrations (in mM): 125 NaCl, 2.5 KCl, 26 $NaHCO_3$, 1.25 $NaH_2PO_4$, 25 glucose, 2 $CaCl_2$, and 1 $MgCl_2$, with an osmolarity of 310 mOs/kg. This solution was bubbled with a gas mixture of 95% $O_2$ and 5% $CO_2$ and maintained at a temperature range of 32−35°C. Additionally, the aCSF contained 100 µM picrotoxin (Cat. No.1128, Tocris Bioscience, Bristol, UK) to inhibit $GABA_A$ receptor-mediated inhibitory postsynaptic currents. Micropipettes (BF150-86-10; Sutter Instrument, Novato, CA) were fabricated using a P1000 puller (Sutter Instrument) and subsequently fire-polished with a microforge. For voltage-clamp recordings, the pipette resistance was maintained between 4 and 5 megohms, while for current-clamp recordings, the resistance ranged from 5 to 7 megohms. The intracellular solution was a potassium-based internal solution with the following composition (in mM): 130 K-gluconate, 10 KCl, 10 HEPES, 0.1 EGTA, 10 $Na_2$-phosphocreatine, 4 Mg-ATP, 0.3 $Na_2$-GTP, and 0.05 Alexa Fluor 488, adjusted to a pH of 7.25 and supplemented with 0.4% biocytin (B4261, Sigma-Aldrich, St. Louis, MO, USA). During voltage-clamp recordings (EPSC recordings), the membrane potential was held at −80 mV. Series resistance (Rs) was monitored for stability, and series resistance compensation was not applied. The bridge balance and pipette capacitance neutralization were meticulously adjusted and verified for stability during current-clamp recordings. Voltage traces were filtered at 10 kHz and digitized at 50 kHz, while current traces were filtered at 2 kHz and digitized at 20 kHz for current-clamp recordings. Data acquisition was performed using AxoGraph X software (AxoGraph, Sydney, Australia) interfaced with an NI USB-6363 device (National Instruments, Austin, TX). The liquid junction potential was corrected by −10 mV.

**Two-photon glutamate uncaging**

The imaging and single-spine stimulation using two-photon microscopy have been described previously (*44*). Briefly, neurons were visualized during whole-cell patch-clamp recordings using Alexa Fluor 488, excited by a two-photon laser at 950 nm (MaiTai DeepSee: Specra-Physics, Milpitas, CA). A glass micropipette containing 10 mM MNI-caged L-glutamate (MNI-Glu, Cat. No. 1490, Tocris Bioscience, Bristol, UK) with a tip diameter of approximately 100 µm was placed roughly 50–100 µm from the surface of the targeted dendritic area of the slice. The MNI-Glu was applied to the targeted dendritic area of the slice using the pressure-puff method. Immediately, uncaging was performed by a two-photon laser at 720 nm directly on the selected spine, which photolyzed MNI-Glu and released active glutamate. The stimulation laser was precisely aligned above the spine to ensure that glutamate activation was localized, avoiding stimulation of nearby structures.

**Confocal imaging**

Confocal fluorescence imaging was executed using an FV4000 confocal microscope (Evident Scientific, Tokyo, Japan). The imaging utilized a UPlanXApo 20× objective lens with a numerical aperture of 0.80, and a digital zoom factor of 1× was applied. Images were captured at a resolution of 1024 × 1024 pixels, with a scanning rate of 2 ms per pixel. Z-stack images were collected at 0.5 μm intervals along the z-axis. A laser with a 488 nm excitation wavelength was employed for detecting fluorescence. All imaging parameters were consistently applied across all samples.

**Immunohistochemistry**
To visualize the neurons that were recorded, biocytin was added to the internal pipette solution during whole-cell patch clamp recordings. After completing the recordings, brain slices were fixed in 4% paraformaldehyde in phosphate-buffered saline (PBS) at 4°C overnight. The fixed slices were then thoroughly washed with PBS and incubated with streptavidin conjugated to Alexa Fluor 488 (1:1000; S32354, Invitrogen, Waltham, MA, USA) at 4°C overnight to label the biocytin-filled cells. Following the washing process, the slices were mounted using an anti-fade mounting medium and covered with a coverslip.

**Pathological examination**
Histopathological examination was conducted on formalin-fixed, paraffin-embedded human brain samples after whole-cell patch clamp recordings. The brain tissues were cut into 6-μm-thick sections. The sections were stained with hematoxylin and eosin (HE). For immunohistochemical studies, a monoclonal antibody raised against Ki67 (M7240, 1:50, Dako, Agilent Technologies Japan, Ltd. Tokyo, Japan) was used. The sections were processed using a Ventana Discovery automated immunostainer (Roche, Basel, Switzerland) (*45*). All stained sections were assessed by a board-certified neuropathologist using a light microscope (ECLIPS Ni, Nikon, Tokyo, Japan) in order to evaluate cancer cell infiltration as well as any neuropath logic abnormalities.

**Quantification and statistical analysis**
Voltage and current waveform analysis was conducted using AxoGraph X and Igor Pro 9.05 (WaveMetrics Inc., Lake Oswego, OR). Input resistance was calculated from the voltage responses to hyperpolarizing current steps (duration 1000 ms), measuring the response at the end of the step. Voltage sag ratio was calculated as (peak-steady state)/peak for current injections of −120 pA. The AP voltage threshold was defined by the voltage when the first time derivative exceeded 20 V/s. The voltage threshold was defined as the EPSP amplitude when an AP was fired. Possible interneurons judged by their characteristic passive and firing properties (*46*) were discarded in this study. Spontaneous EPSCs (sEPSCs) were analyzed using AxoGraph X based on a template-matching algorithm method (*47*). Amplitudes of EPSC > 4 times the standard deviation of the recording noise were detected. Rs was monitored for stability, and data were just used from cells with Rs <20 MΩ. Data were only used EPSCs with rise-time <10 ms, decay time constant <50 ms and half-width <50 ms. Similarly, spontaneous EPSPs (sEPSPs) were analyzed also using the template-matching algorithm. Amplitudes of EPSP > 6 times the standard deviation of the recording noise were detected since overlapped EPSPs which is attributable to the longer time course of sEPSP impedes accurate analysis. Membrane resistance (Rm) is also critical for the time course of EPSCs therefore Rm was monitored for stability carefully during recordings, and data were just used from cells with Rs ranged from 80 to 200 MΩ. Data were only used EPSPs with rise-time <20 ms, decay time constant <100 ms and half-width <100 ms. For sEPSP recordings, the membrane potential was kept at −70 mV using a subtle current injection basically, but events with baseline voltage of EPSPs >−65 mV were excluded to prevent from contamination of NMDA components which cause long time course events.

**Computational model**

We used a two-compartment neuron model consisting of somatic and dendritic compartments (*23*). The dendritic membrane potential of the output neuron at time $t$ was computed as:

$$v(t) = \mathbf{w} \cdot \boldsymbol{e}(t),$$

where $\mathbf{w}$ denotes the synaptic weights projecting to the dendrite and $\boldsymbol{e}$ represents the postsynaptic potentials evoked by presynaptic inputs. The initial synaptic weights $\mathbf{w}$ were drawn from a Gaussian distribution with zero mean and a standard deviation of $1/\sqrt{N}$, and their absolute values were used as initial values.

The dynamics of the somatic membrane potential were described by

$$\dot{U}(t) = -g_{\mathrm{L}}U(t) + g_{\mathrm{D}}\big(-U(t) + V(t)\big),$$

where $g_{\mathrm{L}} = 0.2$ and the conductance between the two compartments is $g_{\mathrm{D}} = 1$. The somatic firing rate was calculated with the nonlinear activation function given as:

$$\phi(U) = \big[1 + \exp\big(-\beta(U - \theta)\big)\big]^{-1},$$

where $\beta$ and $\theta$ were defined as follows:

$$\beta = \sigma(t)^{-1}\beta_0,$$

$$\theta = \mu(t) + \sigma(t)\theta_0.$$

Here, $\mu(t)$ and $\sigma(t)$ are the mean and variance of the membrane potential, respectively, over a sufficiently long period of 500 ms (*23*). We set $\beta_0 = 5$ and $\theta_0 = 1$ throughout this study. The dynamic activation function is important for avoiding a trivial solution of the learning rule described below as shown previously (*23*). We solved differential equations numerically using the Euler method with an integration time step of 0.1 ms throughout this study. Species-specific parameters were set as follows: for the human-type neuron, $\tau = 4.8$ and $\tau_s = 0.8$; for the mouse-type neuron, $\tau = 7.3$ and $\tau_s = 1.2$; and for the macaque-type neuron, $\tau = 7.5$ and $\tau_s = 1.2$.

Afferent inputs were modeled as Poisson spike trains of $N_{\mathrm{in}}$ input neurons:

$$X_k^{\mathrm{ext}}(t) = \sum_q \delta\big(t - t_{k,q}^{\mathrm{ext}}\big),$$

where $\delta$ is the Dirac's delta function and $t_{k,q}^{\mathrm{ext}}$ is the time of the $q$-th spike generated by the $k$-th input neuron. We used $N_{\mathrm{in}} = 2{,}000$ input neurons, each generating Poisson spikes at a constant mean rate of 5 Hz. The postsynaptic potential evoked by the $k$-th input was calculated as

$$\tau_s \dot{I}_k^{\mathrm{ext}} = -I_k^{\mathrm{ext}} + \frac{1}{\tau}X_k^{\mathrm{ext}},$$

$$\dot{e}_k^{\mathrm{ext}} = -\frac{e_k^{\mathrm{ext}}}{\tau} + e_0 I_k^{\mathrm{ext}},$$

where $\tau_s$ is a synaptic time constant and $e_0$ is a scale of the postsynaptic potential defined as:

$$e_0 = \frac{\tau - \tau_s}{\left(\frac{\tau}{\tau_s}\right)^{-\frac{\tau_s}{\tau-\tau_s}} - \left(\frac{\tau}{\tau_s}\right)^{-\frac{\tau}{\tau-\tau_s}}}.$$

We used the predictive learning rule proposed previously to train the synaptic weight (*23, 26*):

$$\Delta\mathbf{w} = \varepsilon[f(t) - \phi_0(V^*)]\boldsymbol{e},$$

where $\varepsilon$ is the learning rate of $5 \times 10^{-5}$ and $\phi_0$ is the sigmoidal function:

$$\phi_0(x) = \big[1 + \exp\big(-\beta_0(x - \theta_0)\big)\big]^{-1},$$

and $V^*(t) = \frac{g_D}{g_D + g_L}V(t)$ is the dendritic prediction of somatic firing rate (*23*). The time-varying signal $f(t)$ serves as a teaching signal for dendritic synapses. In the unsupervised learning regime, $f(t) = \phi(U)$ (*23*), whereas in the supervised regime, $f(t) = 1$ for target patterns and $f(t) = 0$ otherwise. To restrict all connections to be excitatory, synaptic weights were clipped to non-negative values at every time step. The prediction error in the unsupervised regime (Fig. 5D) was calculated as the mean squared error (MSE) between the somatic and dendritic firing rates.

In the analyses shown in Fig. 5E–I, input spike patterns were randomly omitted according to the designated probabilities, while the overall input firing rate was kept constant at 5 Hz across conditions. Success rate was defined as the probability of reaching 50% of the maximum firing rate within 3 ms after pattern onset. Error bars in all simulations were calculated from 10 independent runs.

**Statics**

See Table 2 for the details. Throughout the text and figure legends, all data are presented as mean ± s.e.m. Given that the biological datasets in this study frequently did not conform to a normal distribution, non-parametric statistical methods were employed for group comparisons. The Mann-Whitney U test was utilized for two-group comparisons, the Kruskal-Wallis test for multiple comparisons, and the Steel test for post-hoc analyses. Datasets depicted as cumulative distribution functions were compared using the Kolmogorov-Smirnov test. Statistical analyses were conducted using R or EZR (*48*).

| Patient No. | Sex | Age | Resection location | Diagnosis | Pathological examination |
|---|---|---|---|---|---|
| 1 | Female | 72 | Parietal lobe | Metastatic brain tumor | No cancer cell infiltration |
| 2 | Male | 46 | Frontal lobe | Glioma | No cancer cell infiltration |
| 3 | Male | 54 | Frontal lobe | Glioma | No cancer cell infiltration |
| 4 | Female | 57 | Frontal lobe | Metastatic brain tumor | No cancer cell infiltration |
| 5 | Male | 32 | Temporal lobe | Epilepsy | No abnormal findings |
| 6 | Male | 81 | Temporal lobe | Glioma | Cancer cell infiltration |

**Table 1. Demographic data.**
Patient #6 was excluded from the data analysis after a thorough pathological inspection.

| | sample | | statistics | | | | | |
|---|---|---|---|---|---|---|---|---|
| | a description | size (n) | methods | comparison | p values / correlation coefficient / coefficient of determination | | | |
| **Figure 1C-L** | Acute slice, prefrontal or parietal cotex | Mouse = 12 neurons / 4 mice<br>Rats = 16 neurons / 3 rats<br>Marmoset = 9 neurons / 4 marmosets<br>Macaque = 14 neurons / 3 macaques<br>Human = 28 neurons / 5 humans | Kruskal-Wallis test (post-hoc: Steel test) | | Resting membrane potential | Membrane resistance | Membrane time constant | AP threshold |
| | | | | Human vs Mouse | 0.43422 | 0.30008 | 0.00139 | 0.65223 |
| | | | | Human vs Rat | 0.68969 | 0.98956 | 0.00159 | 0.98828 |
| | | | | Human vs Marmoset | 0.61467 | 0.29102 | 0.00004 | 0.41730 |
| | | | | Human vs Macaque | 0.88470 | 0.33073 | 0.42995 | 0.96710 |
| | | | | | AP height | AP peak | AP rise time | AP half width |
| | | | | Human vs Mouse | 0.30012 | 0.60995 | 0.01621 | 0.40538 |
| | | | | Human vs Rat | 0.37004 | 0.20555 | 0.98690 | 0.00009 |
| | | | | Human vs Marmoset | 0.48885 | 0.99900 | 0.99279 | 0.02211 |
| | | | | Human vs Macaque | 1.00000 | 0.71085 | 0.99333 | 0.00406 |
| | | | | | Rheobase | Sag | | |
| | | | | Human vs Mouse | 0.38567 | 0.00252 | | |
| | | | | Human vs Rat | 0.99595 | 0.03548 | | |
| | | | | Human vs Marmoset | 0.01355 | 0.13576 | | |
| | | | | Human vs Macaque | 0.98215 | 0.71972 | | |
| **Figure 2B-D** | Acute slice, prefrontal or parietal cotex | Mouse = 20 neurons / 8 mice<br>Rats = 11 neurons / 5 rats<br>Marmoset = 16 neurons / 6 marmosets<br>Macaque = 7 neurons / 3 macaques<br>Human = 25 neurons / 5 humans | Kruskal-Wallis test (post-hoc: Steel test) | | Rise time | Decay time | Amplitude | |
| | | | | Human vs Mouse | 0.00000 | 0.00000 | 0.00242 | |
| | | | | Human vs Rat | 0.00003 | 0.00014 | 0.01146 | |
| | | | | Human vs Marmoset | 0.00000 | 0.00000 | 0.80915 | |
| | | | | Human vs Macaque | 0.00045 | 0.00054 | 0.99991 | |
| **Figure 2E-G** | Acute slice, prefrontal or parietal cotex | Mouse = 20 neurons / 8 mice<br>Rats = 11 neurons / 5 rats<br>Marmoset = 16 neurons / 6 marmosets<br>Macaque = 7 neurons / 3 macaques<br>Human = 25 neurons / 5 humans | Two-sample Kolmogorov-Smirnov test | | Rise time | Decay time | Amplitude | |
| | | | | Human vs Mouse | 0.00000 | 0.00000 | 0.00000 | |
| | | | | Human vs Rat | 0.00000 | 0.00000 | 0.00000 | |
| | | | | Human vs Marmoset | 0.00000 | 0.00000 | 0.44167 | |
| | | | | Human vs Macaque | 0.00000 | 0.00000 | 0.16097 | |
| **Figure 2H-J** | Acute slice, prefrontal or parietal cotex | Rodent = 31 neurons / 13 animals<br>Non-human primate = 23 neurons / 9 animals<br>Human = 25 neurons / 5 humans | Two-sample Kolmogorov-Smirnov test | | Rise time | Decay time | Amplitude | |
| | | | | Human vs Rodent | 0.00000 | 0.00000 | 0.00000 | |
| | | | | Human vs Non-human primate | 0.00000 | 0.00000 | 0.34703 | |
| **Figure 3B-C** | Acute slice, prefrontal or parietal cotex | Mouse = 14 neurons / 6 mice<br>Rats = 12 neurons / 4 rats<br>Marmoset = 11 neurons / 7 marmosets<br>Macaque = 6 neurons / 2 macaques<br>Human = 13 neurons / 2 humans | Kruskal-Wallis test (post-hoc: Steel test) | | Rise time | Decay time | | |
| | | | | Human vs Mouse | 0.00442 | 0.00062 | | |
| | | | | Human vs Rat | 0.00505 | 0.02045 | | |
| | | | | Human vs Marmoset | 0.00117 | 0.00041 | | |
| | | | | Human vs Macaque | 0.03935 | 0.04986 | | |
| **Figure 3D-E** | Acute slice, prefrontal or parietal cotex | Mouse = 14 neurons / 6 mice<br>Rats = 12 neurons / 4 rats<br>Marmoset = 11 neurons / 7 marmosets<br>Macaque = 6 neurons / 2 macaques<br>Human = 13 neurons / 2 humans | Two-sample Kolmogorov-Smirnov test | | Rise time | Decay time | | |
| | | | | Human vs Mouse | 0.00019 | 0.00041 | | |
| | | | | Human vs Rat | 0.00000 | 0.00364 | | |
| | | | | Human vs Marmoset | 0.00000 | 0.00012 | | |
| | | | | Human vs Macaque | 0.00003 | 0.00012 | | |
| **Figure 3F-G** | Acute slice, prefrontal or parietal cotex | Rodent = 26 neurons / 10 animals<br>Non-human primate = 17 neurons / 9 animals<br>Human = 13 neurons / 2 humans | Two-sample Kolmogorov-Smirnov test | | Rise time | Decay time | | |
| | | | | Human vs Rodent | 0.00003 | 0.00020 | | |

| | | | | | | | | | |
|---|---|---|---|---|---|---|---|---|---|
| | | | | Human vs Non-human primate | 0.00000 | 0.00096 | | | |
| **Figure 4C-D** | Acute slice, prefrontal or parietal cotex | Mouse = 30 spines / 3 neurons / 1 mouse<br>Rats = 15 spines / 4 neurons / 3 rats<br>Marmoset = 16 spines / 5 neurons / 3 marmosets<br>Macaque = 13 spines / 1 neuron / 1 macaque<br>Human =14 spines / 2 neurons / 1 human | Kruskal-Wallis test (post-hoc: Steel test) | | Rise time | Decay time | | | |
| | | | | Human vs Mouse | 0.00505 | 0.00308 | | | |
| | | | | Human vs Rat | 0.00015 | 0.00132 | | | |
| | | | | Human vs Marmoset | 0.00014 | 0.00264 | | | |
| | | | | Human vs Macaque | 0.00266 | 0.27651 | | | |
| **Figure 5F** | Model neurons, unsupervised learning | Mouse = 10 neurons<br>Macaque = 10 neurons<br>Human = 10 neurons | Kruskal-Wallis test (post-hoc: Steel test) | | Success rate | | | | |
| | | | | Human vs Mouse | 0.01207 | | | | |
| | | | | Human vs Macaque | 0.00305 | | | | |
| **Figure 5I** | Model neurons, supervised learning | Mouse = 10 neurons<br>Macaque = 10 neurons<br>Human = 10 neurons | Kruskal-Wallis test | | Success rate | | | | |
| | | | | Human vs Mouse | n.s. | | | | |
| | | | | Human vs Macaque | n.s. | | | | |
| **Figure S2A-E** | Acute slice, prefrontal or parietal cotex | Mouse = 20 neurons / 8 mice<br>Rats = 11 neurons / 5 rats<br>Marmoset = 4 neurons / 2 marmosets<br>Macaque = 7 neurons / 3 macaques<br>Human = 20 neurons / 4 humans | Pearson correlation coefficient | Rise time vs Soma location | Mouse | Rat | Marmoset | Macaque | Human |
| | | | | | 0.30350 | 0.04333 | 0.62690 | 0.51860 | 0.21020 |
| | | | | | Mouse | Rat | Marmoset | Macaque | Human |
| | | | | | 0.24221 | 0.61660 | -0.37311 | -0.29639 | 0.29283 |
| **Figure S2F-J** | Acute slice, prefrontal or parietal cotex | Mouse = 20 neurons / 8 mice<br>Rats = 11 neurons / 5 rats<br>Marmoset = 4 neurons / 2 marmosets<br>Macaque = 7 neurons / 3 macaques<br>Human = 20 neurons / 4 humans | Pearson correlation coefficient | Decay time vs Soma location | Mouse | Rat | Marmoset | Macaque | Human |
| | | | | | 0.35930 | 0.01313 | 0.90270 | 0.26000 | 0.07035 |
| | | | | | Mouse | Rat | Marmoset | Macaque | Human |
| | | | | | 0.21649 | 0.71642 | -0.09735 | -0.49381 | 0.41295 |
| **Figure S3A** | Acute slice, prefrontal or parietal cotex | Perfusion = 11 neurons / 6 animals<br>Non-perfusion = 9 neurons / 2 animals | Mann-Whitney U test | Perfusion vs Non-perfusion | Rise time | Decay time | | | |
| | | | | | 0.60270 | 0.65560 | | | |
| **Figure S3B** | Acute slice, prefrontal or parietal cotex | Perfusion = 5 neurons / 2 animals<br>Non-perfusion = 6 neurons / 3 animals | Mann-Whitney U test | Perfusion vs Non-perfusion | Rise time | Decay time | | | |
| | | | | | 0.53680 | 0.32900 | | | |
| **Figure S3C** | Acute slice, prefrontal or parietal cotex | Perfusion = 11 neurons / 4 animals<br>Non-perfusion = 5 neurons / 2 animals | Mann-Whitney U test | Perfusion vs Non-perfusion | Rise time | Decay time | | | |
| | | | | | 0.00870 | 0.01328 | | | |
| **Figure S3D** | Acute slice, prefrontal or parietal cotex | Perfusion = 5 neurons / 2 animals<br>Non-perfusion = 2 neurons / 1 animal | Mann-Whitney U test | Perfusion vs Non-perfusion | Rise time | Decay time | | | |
| | | | | | 1.00000 | 0.09524 | | | |
| **Figure S4A-E** | Acute slice, prefrontal or parietal cotex | Mouse = 30 spines / 3 neurons / 1 mouse<br>Rats = 15 spines / 4 neurons / 3 rats<br>Marmoset = 16 spines / 5 neurons / 3 marmosets<br>Macaque = 13 spines / 1 neuron / 1 macaque<br>Human =14 spines / 2 neurons / 1 human | Pearson correlation coefficient | Distance from soma vs Rise time | Mouse | Rat | Marmoset | Macaque | Human |
| | | | | | 0.02229 | 0.00723 | 0.57710 | 0.50370 | 0.01699 |
| | | | | | Mouse | Rat | Marmoset | Macaque | Human |
| | | | | | 0.41583 | 0.66157 | 0.15084 | 0.20405 | -0.62438 |
| **Figure S4F-J** | Acute slice, prefrontal or parietal cotex | Mouse = 30 spines / 3 neurons / 1 mouse<br>Rats = 15 spines / 4 neurons / 3 rats<br>Marmoset = 16 spines / 5 neurons / 3 marmosets<br>Macaque = 13 spines / 1 neuron / 1 macaque<br>Human =14 spines / 2 neurons / 1 human | Pearson correlation coefficient | Distance from soma vs Decay time | Mouse | Rat | Marmoset | Macaque | Human |
| | | | | | 0.17460 | 0.90850 | 0.70210 | 0.38860 | 0.85890 |
| | | | | | Mouse | Rat | Marmoset | Macaque | Human |
| | | | | | -0.25458 | -0.03249 | -0.10379 | 0.26126 | 0.05234 |
| **Figure S5A-E** | Acute slice, prefrontal or parietal cotex | Mouse = 20 neurons / 8 mice<br>Rats = 11 neurons / 5 rats<br>Marmoset = 16 neurons / 6 marmosets<br>Macaque = 7 neurons / 3 macaques<br>Human = 25 neurons / 5 humans | Pearson correlation coefficient | Age vs Rise time | Mouse | Rat | Marmoset | Macaque | Human |
| | | | | | 0.62910 | 0.10390 | 0.02827 | 0.74940 | 0.00788 |
| | | | | | Mouse | Rat | Marmoset | Macaque | Human |
| | | | | | -0.11505 | 0.51643 | -0.54716 | 0.14930 | -0.51881 |
| **Figure S5F-J** | Acute slice, prefrontal or parietal cotex | Mouse = 20 neurons / 8 mice<br>Rats = 11 neurons / 5 rats<br>Marmoset = 16 neurons / 6 marmosets<br>Macaque = 7 neurons / 3 macaques<br>Human = 25 neurons / 5 humans | Pearson correlation coefficient | Age vs Decay time | Mouse | Rat | Marmoset | Macaque | Human |
| | | | | | 0.61120 | 0.03234 | 0.00776 | 0.11900 | 0.00805 |
| | | | | | Mouse | Rat | Marmoset | Macaque | Human |
| | | | | | -0.12104 | 0.64439 | -0.63854 | 0.64343 | -0.51762 |

**Table 2. Statical details.**
The statistics used, the sample size and the sample type were described.

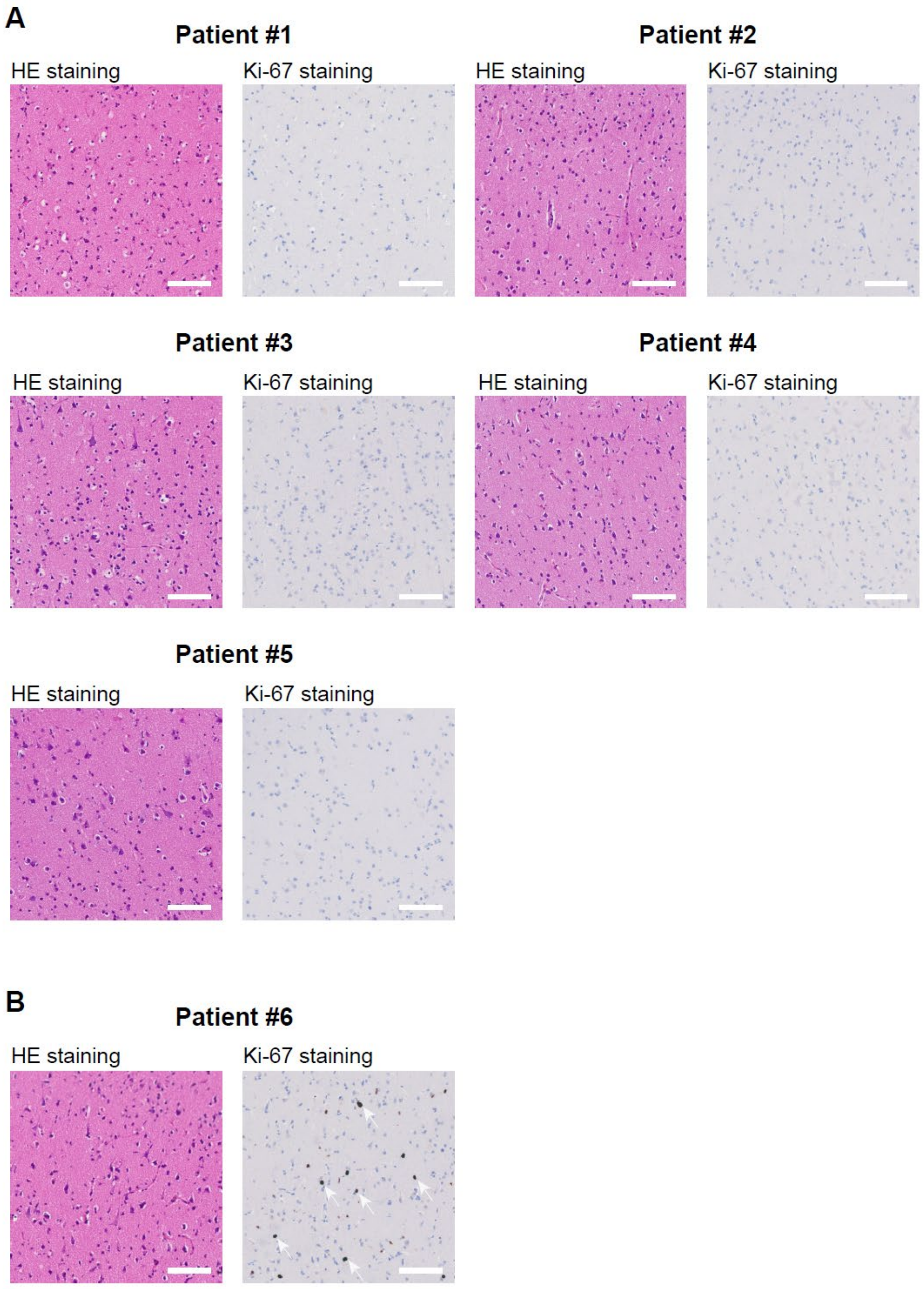


**Supplementary Figure 1.**
(**A**) A pathological diagnosis was made using Hematoxylin and Eosin (HE) staining and anti-Ki67 staining of brain specimens that were judged to have no cancer cell infiltration. (**B**) HE and anti-Ki67 staining of brain specimens excluded from this study due to pathological evidence of tumor infiltration. The arrows indicate Ki-67-positive cells.

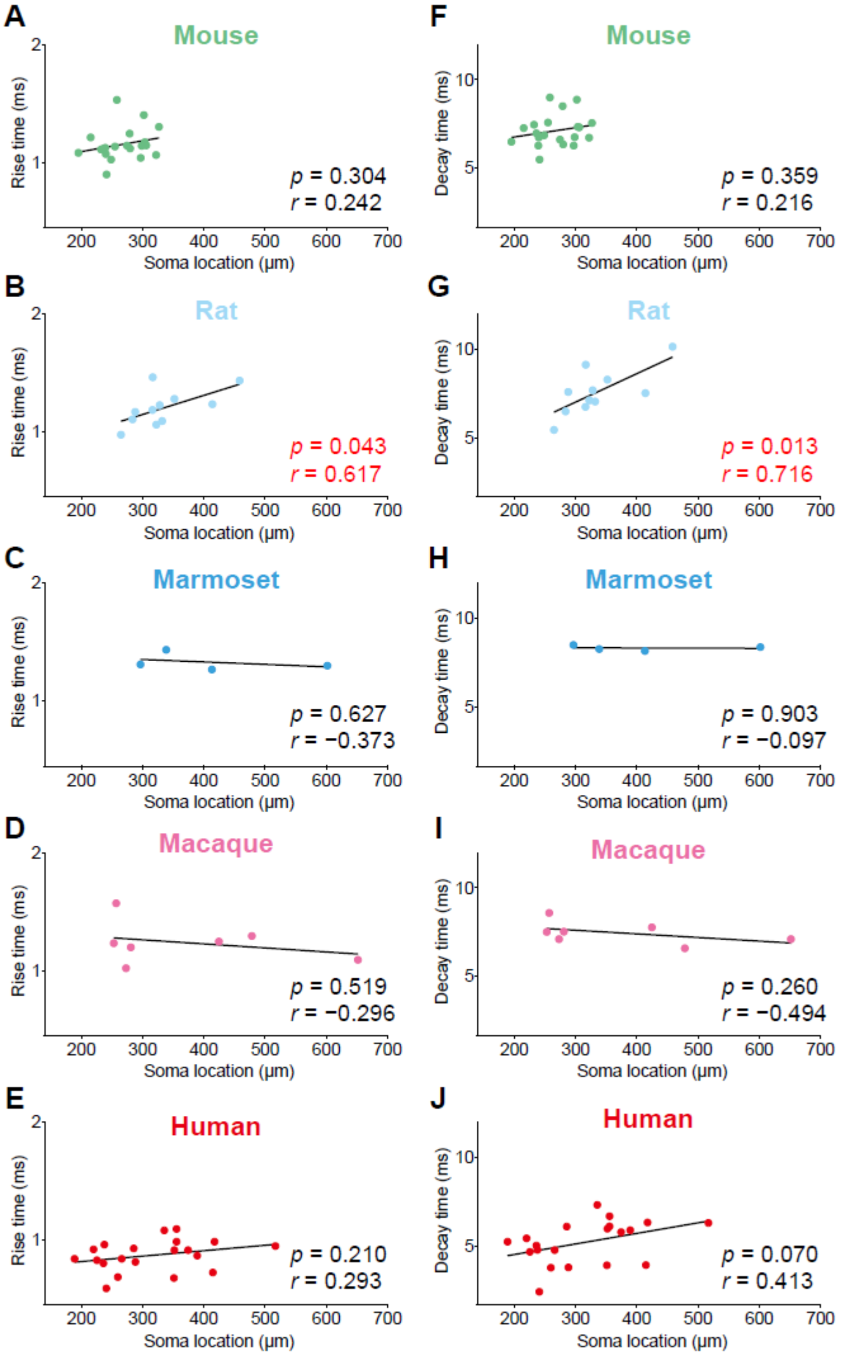


**Supplementary Figure 2. The location of the recorded cells in the cortex and their relationship to the temporal kinetics of EPSCs.**
Graphs showing the distance from the pia mater to neuronal cell bodies, as well as its relationship to the rise and decay times of EPSCs. Pearson's correlation coefficient.

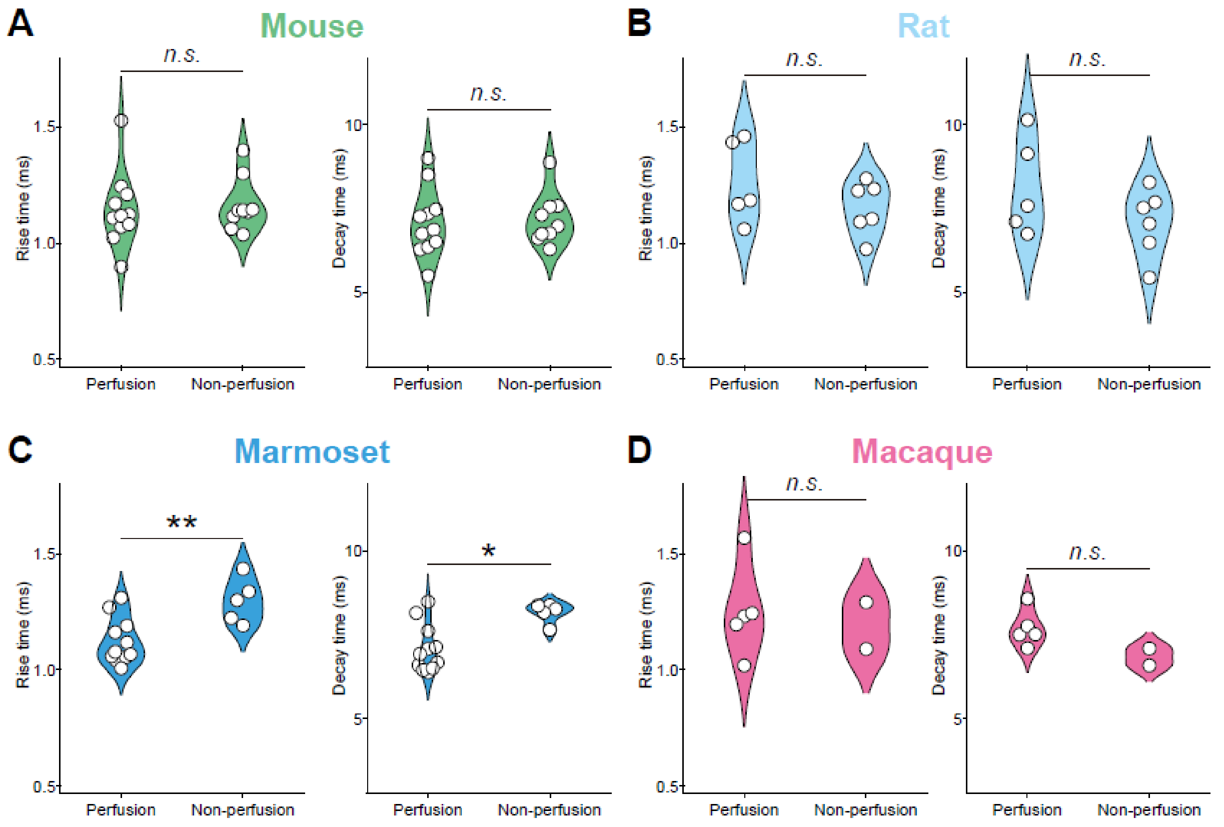


**Supplementary figure 3. The influence of cardiac perfusion prior to brain resection on the time course of the EPSC.**
(**A–D**) The standard protocol for preparing acute brain slices from animal models involves transcardiac perfusion with an ice-cold physiological solution prior to resection of the brain. As such procedures are not performed on human patients, however, the effect of perfusion was evaluated in animal models. Mouse (**A**, Perfusion $n$ = 11, Non-perfusion $n$ = 9; Rise time, $P$ = 0.603, Decay time, $P$ = 0.656). Rat (**B**, Perfusion $n$ = 5, Non-perfusion $n$ =6; Rise time, $P$ = 0.537, Decay time, $P$ = 0.329). Marmoset (**C**, Perfusion $n$ = 11, Non-perfusion $n$ = 5; Rise time $P$ = 0.009, Decay time $P$ = 0.013). Macaque (**D**, Perfusion $n$ = 5, Non-perfusion $n$ = 2; Rise time, $P$ = 1, Decay time, $P$ = 0.095). Mann-Whitney U test. $*P < 0.05$, $**P < 0.01$.

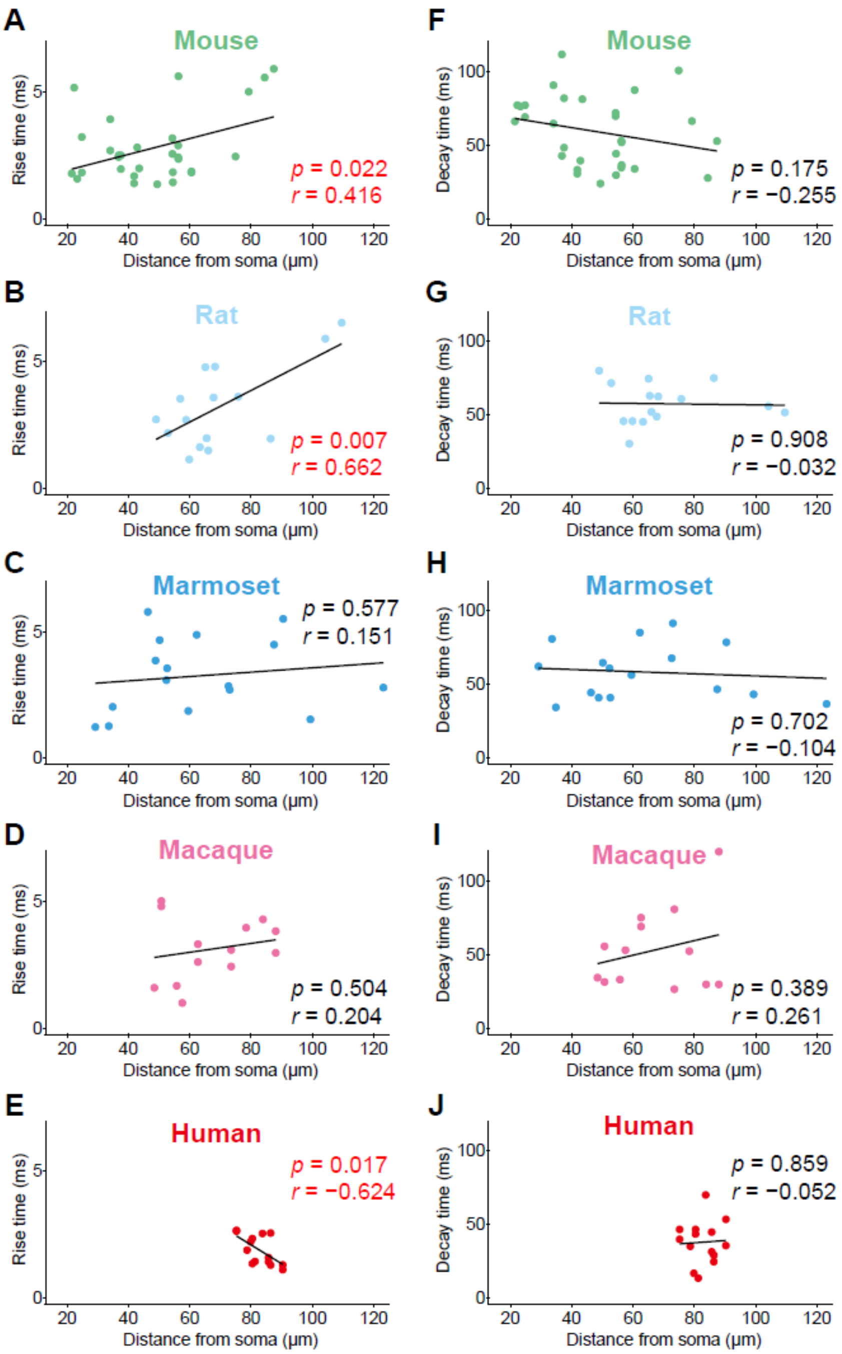


**Supplementary figure 4. No consistent effect of the synapse-to-soma distance on the EPSP kinetics.**

(**A–E**) The distance of the synapse from the soma, as well as the corresponding uncaging-evoked EPSP kinetics, were precisely measured using single-spine stimulation and two-photon glutamate uncaging. Mouse (**A, F**), Rat (**B, G**), Marmoset (**C, H**), Macaque (**D, I**), Human (**E, J**). Pearson's correlation coefficient.

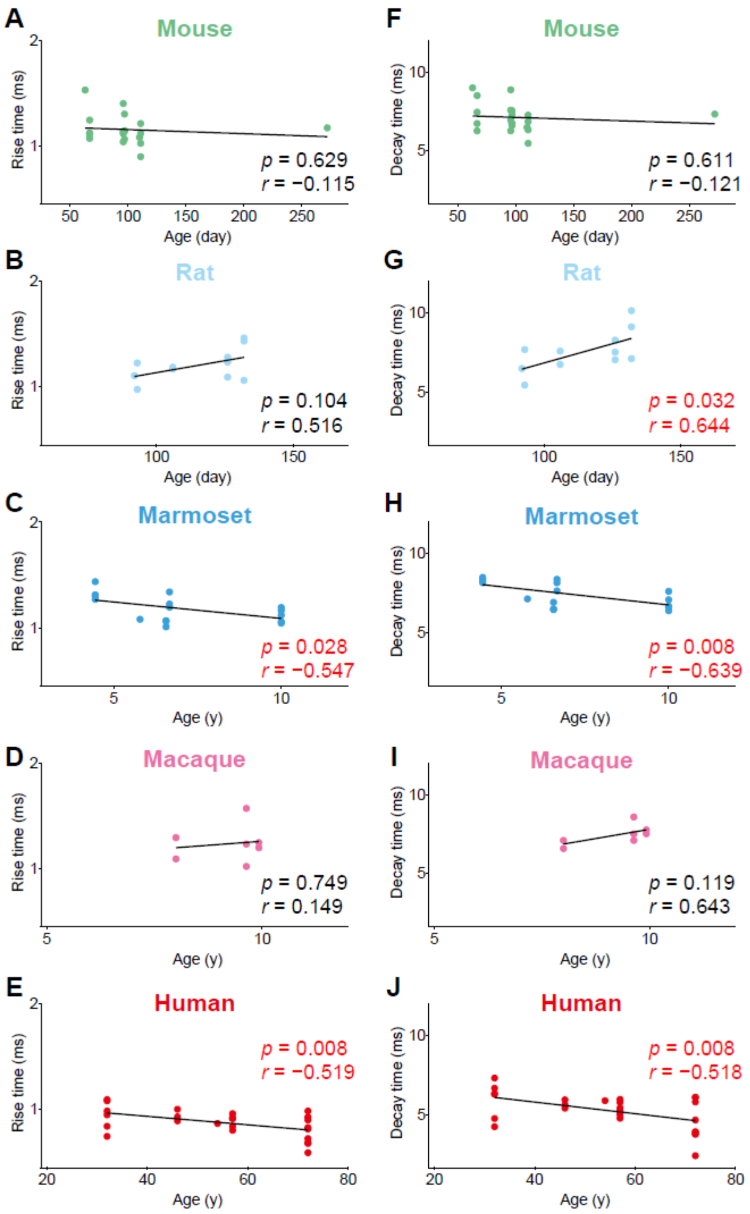


**Supplementary figure 5. The relationship between donor age and the EPSCs kinetics.** (**A–E**) The correlation between the donor age and the rise time of EPSCs. Mouse (**A, F**), Rat (**B, G**), Marmoset (**C, H**), Macaque (**D, I**), Human (**E, J**). Pearson's correlation coefficient.
